# Efficient transformation and genome editing of watermelon assisted by genes that encode developmental regulators

**DOI:** 10.1101/2021.11.05.467370

**Authors:** Wenbo Pan, Zhentao Cheng, Zhiguo Han, Hong Yang, Wanggen Zhang, Huawei Zhang

**Author notes:** These authors contributed equally: Wenbo Pan and Zhentao Cheng. These authors contributed equally: Huawei Zhang and Wanggen Zhang.

## Abstract

The Cucurbitaceae contains multiple species of important food plants. But most of them are difficult to be genetically transformed. Watermelon is one of the most important fruit species of Cucurbitaceae, and it is a model horticulture crops. Its low transgenic efficiency is the major bottleneck in functional genome research and genome editing-based breeding. Here, with the help of genes that encode developmental regulators (DRs), in particular *AtGRF5*, and an appropriate *Agrobacterium* strain (GV3101), we have significantly increased the transformation efficiency of watermelon to about 25%, which is a 40-fold increase compared with a traditional vector. *AtGRF5-*mediated transformation increased the number of transformable watermelon genotypes without causing obvious side effects. Moreover, when applying this strategy to CRISPR/Cas9-mediated genome editing, *clpds* knockout mutants were generated in the T0 generation. Our results show that *AtGRF5* is a powerful and robust tool to effectively create transgenic plants or knockout mutations in watermelon. Similar strategies using DRs might also be able to overcome the transformation barriers in many other Cucurbitaceae species.

## Introduction

The Cucurbitaceae is an important family of flowering plants containing multiple species of important food plants such as melons, cucumbers, squashes, and pumpkins. However, a highly efficient genetic transformation system has not been established for most of these species (Guo *et al*., 2020, Nanasato and Tabei, 2020). Watermelon (*Citrullus lanatus*) is an economically important fruit crop that is cultivated worldwide. It is a model crop for fruit quality research because of its rich diversity in fruit size, shape, flavor, aroma, texture, peel and flesh color, and nutritional composition (Guo *et al*., 2019). The first high-quality draft genome sequence of watermelon was reported in 2013 (Guo *et al*., 2013). Through pan-genome sequencing, many candidate loci associated with fruit quality traits have since been identified (Guo *et al*., 2019). However, few of these loci have been validated. The major barrier is the low transformation efficiency of watermelon, with only few successful cases of transformation reported (Yu *et al*., 2011, Tian *et al*., 2017, Wang *et al*., 2021a, Wang *et al*., 2021b). For example, in one study only 16 transgenic lines were obtained from about 960 cotyledon fragments (Tian *et al*., 2017), g giving a transformation efficiency of 1.67%. Therefore, methods that improve the transformation efficiency could not only facilitate functional genomic studies in watermelon as well as other horticulture species, but also speed up transgenic breeding and genome editing breeding, which both require genetic transformation.

Reprograming of the cell fate of explants in tissue culture is regulated by genes related to plant development. Plants are well-known for their natural ability to regenerate new tissues or plants from wounded tissues. Previous studies have identified numerous genes that can promote or reprogram cell fate, and they are known as developmental regulators (DRs) (Mendez-Hernandez *et al*., 2019). Several DRs have been reported to improve the regeneration efficiency of various plant species in tissue culture (Mendez-Hernandez *et al*., 2019, Zhang *et al*., 2021). For example, overexpression of the AP2/ERF family transcription factor BABY BOOM (BBM), which has diverse functions in plant development, can promote cell proliferation and ectopic embryo formation in the cotyledons and leaves of *Arabidopsis* (Horstman *et al*., 2017). Co-expression of BBM with the shoot apical meristem identity regulator WUSCHEL (WUS) greatly improves the *in vitro* transformation efficiency of various monocot species, including many maize inbred lines, rice, and sorghum (Lowe *et al*., 2016). Ectopic co-expression of *WUS* and the *Agrobacterium* cytokinin biosynthesis gene *isopentenyl transferase* (*ipt*) is sufficient to induce meristem-like growths and new shoots on seedlings of *Nicotiana benthamiana* (Maher *et al*., 2020). However, constitutive overexpression of both *BBM* and *WUS* causes severe growth defects, such as abnormal development of vegetative and reproductive organs and infertility. Therefore, expression of these DRs needs to be restricted or the genes need to be eliminated during or after tissue culture (Lowe *et al*., 2016). The plant-specific transcription factor proteins called GROWTH-REGULATING FACTORs (GRFs) have also been reported to boost the regeneration and genetic transformation efficiency of various crop plants. GRFs form a transcription complex with GRF INTERACTING FACTOR (GIF) to regulate plant growth and development by recruiting chromatin remodeling complexes and providing cues to the primordial cells of vegetative and reproductive organs (Omidbakhshfard *et al*., 2015). Stable expression of *GRF5* from *Arabidopsis* accelerates shoot organogenesis and increases the genetic transformation efficiency of soybean, canola, and sunflower (Kong *et al*., 2020). Similarly, overexpression of a chimeric protein consisting of the wheat GRF4 and rice GIF1 proteins improves the regeneration efficiency and speed of regeneration in wheat, triticale, and rice, and also increases the number of transformable wheat genotypes (Debernardi *et al*., 2020).

Therefore, we tested the effects of these DRs on watermelon transformation. We cloned several DRs that have been reported to boost somatic embryogenesis in other species. We overexpressed these DRs individually and in combination in two watermelon cultivars and evaluated their efficiency. Furthermore, we added the effective DRs to clustered regularly interspaced short palindromic repeats (CRISPR)/CRISPR-associated protein 9 (Cas9) genome editing vectors and obtained transgenic plants with knockout of the phytoene desaturase gene ClPDS1. The approach we developed here can be applied to functional genomic studies and will facilitate the utilization of genome editing in watermelon and potentially other Cucurbitaceae species. We will test this in the future.

## Results

### Establishment of a watermelon transformation system

To improve the transformation efficiency of watermelon, we first established a fast and simple detection system for watermelon transformation. Both the *neomycin phosphotransferase II* (*nptII*) gene, which confers kanamycin (Kan) resistance in transgenic plants, and the *hygromycin phosphotransferase II* (*hptII*) gene, which confers hygromycin (Hyg) resistance, have been used as selectable markers in watermelon tissue culture. Although both Kan and Hyg are aminoglycoside antibiotics that act as protein synthesis inhibitors, Hyg causes more severe damage than Kan during watermelon tissue culture (Park *et al*., 2005), and the time required for tissue culture in Hyg medium is usually longer than required for Kan; thus, *nptII* was chosen as the selectable marker. In addition, given the high frequency of false positive plants obtained after Kan selection (Park *et al*., 2005), the DsRed2 fluorescent protein, which can be easily observed using the LUYOR-3415RG hand-held lamp (Luyor Corporation, Shanghai, China), was introduced into the binary vector as another marker to visually identify stable transgenic events **(Figure 1a)**. The reporter vector was named pW501.

**Fig. 1.**
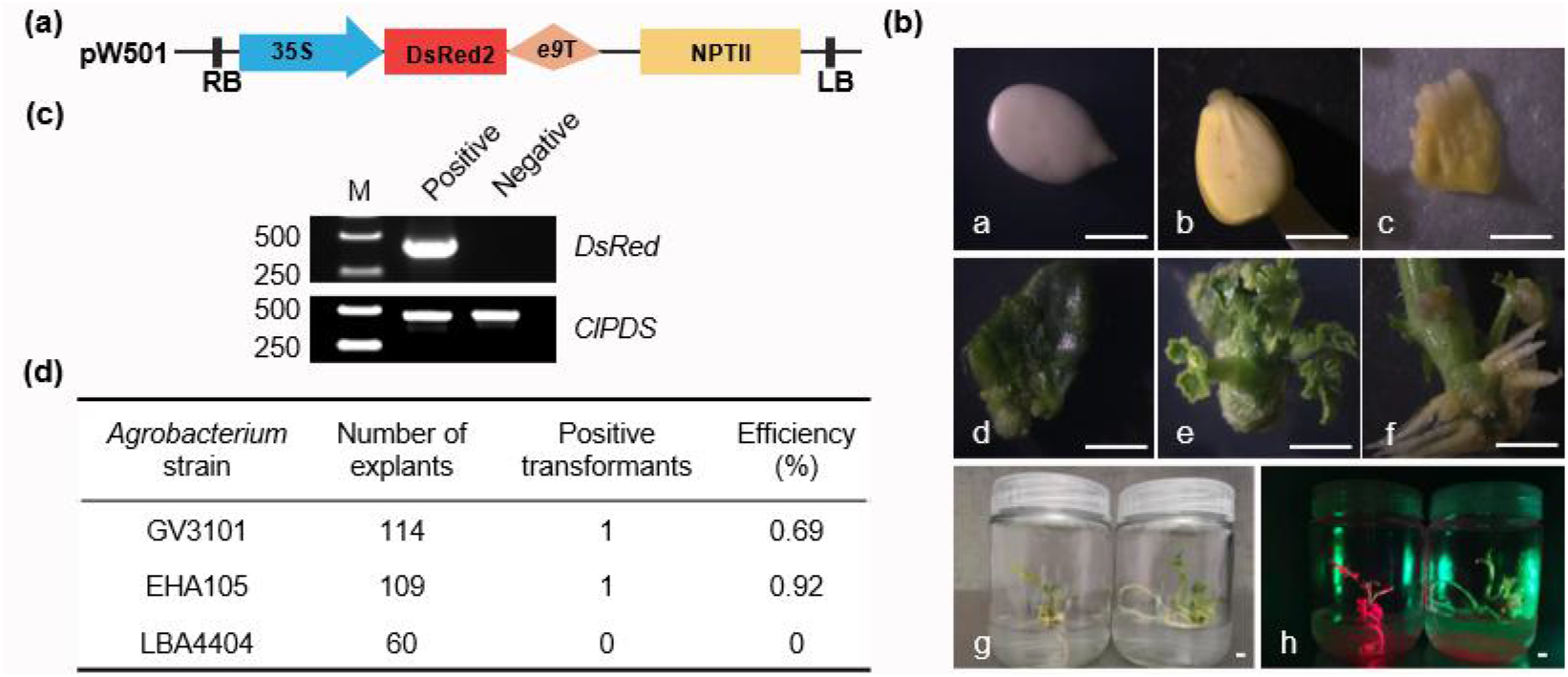
Establishment of a watermelon transformation system. **a**. Schematic diagrams of the reporter vector constructs used for watermelon transformation. **b**. Genetic transformation of the watermelon cultivar WW150. a, watermelon seeds after Sterilization; b, Four-day-old watermelon seedlings; c, cotyledon fragments after co-cultivation; d, callus formed on the selection medium after 2-weeks; e, adventitious buds on the callus; f, adventitious root formed on the regenerated plants; g, seedlings with roots on the rooting medium. The seedling on the left is a transgenic plant, and the seedling on the right is a non-transgenic plant; h, the same plants as in f with the picture taken under a hand-held florescence detection device. Scale bars = 0.5 cm. **c**. PCR verification of the transgenic lines. M, marker. Left number indicate the size of markers. Positive and negative indicate DsRed florescent positive and negative plants. The PCR product of 424 bp corresponding to the DsRed sequence is shown in the upper panel. ClPDS is used as the internal control. **d**. The transformation efficiency of watermelon cultivar WW150 using different *Agrobacterium tumefaciens* strains.

We first tested genetic transformation of the watermelon cultivar WWl50, the female parental line of the widely grown cultivar ‘Xinong No.8’, which is relatively recalcitrant to transformation. The vectors were introduced into three widely used laboratory *Agrobacterium* strains, EHA105, GV3101 and LBA4404, and then delivered to cotyledonary fragments using the established *Agrobacterium*-mediated transformation method (Tian *et al*., 2017). Callus was induced after co-cultivation and growth on selection medium containing 50 mg/L Kan for 2 weeks **(Fig. 1b)**. After about 70 days of selection, adventitious shoots and roots were regenerated, and positive transformation events could be easily detected by a hand-held fluorescent lamp **(Fig. 1b)**. Transgenic-specific PCR result with primers specific to DsRed also demonstrated that the T-DNA fragment is integrated into the watermelon genome in DsRed fluorescent positive lines **(Fig. 1c)**.

The transformation frequencies were determined by counting the number of calli showing at least one regenerated adventitious bud with a DsRed fluorescent signal out of the total number of inoculated cotyledonary fragments. Among the three *Agrobacterium* strains we tested, EHA105 had the highest transformation efficiency, 0.92%, which was comparable with the efficiency reported in previous studies (Tian *et al*., 2017), while the efficiency was 0.69% for GV3101 and no positive transformation events were detected using LBA4404 **(Fig. 1d)**.

### Effects of DRs on the transformation efficiency of watermelon

We predicted that the low transformation efficiency might be mainly due to the low regeneration efficiency of transgenic cells. Several DRs were reported to boost the regeneration efficiency or transgenic efficiency of various plant species in tissue culture (Lowe *et al*., 2016, Mendez-Hernandez *et al*., 2019, Debernardi *et al*., 2020, Kong *et al*., 2020, Maher *et al*., 2020, Zhang *et al*., 2021). To test the effects of DRs on watermelon transformation, we added several DR genes, individually and in combination, to the binary vector pW501. Expression of all DR genes was driven by the *Arabidopsis UBQ10* promoter, except for *WUS*, which was driven by the *Nos* promoter as previously described (Lowe *et al*., 2016) **(Fig. 2a)**.

**Fig. 2.**
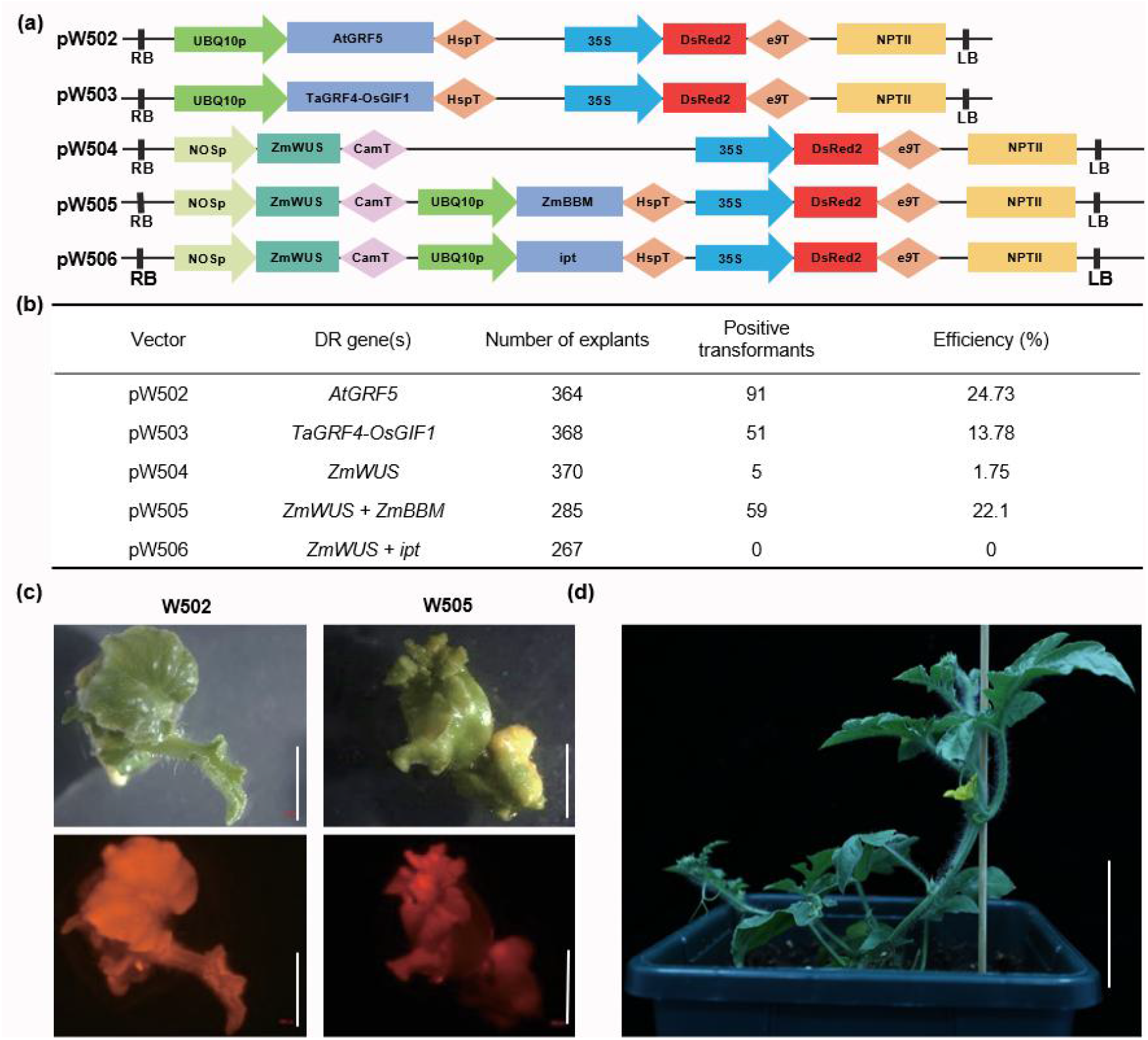
Effects of developmental regulators on watermelon transformation. **a**. Schematic diagrams of constructs with developmental regulators (DRs) used in this study. **b**. The transformation efficiencies of watermelon cultivar WW150 obtained using the indicated constructs. Positive transformation events were defined as calli showing at least one regenerated adventitious bud expressing the DsRed fluorescent signal. **c**. Representative images of adventitious shoots generated by the indicated vectors (upper panels). The fluorescent DsRed2 signals are shown in the lower panels. Scale bars = 0.5 cm. **d**. The growth phenotype of a watermelon plant transformed with the pW502 vector 2 weeks after being transferred to soil. Scale bar = 4 cm.

All of these vectors were introduced into the *Agrobacterium* strain GV3010 and then transformed into the watermelon cultivar WWl50. Some of these DR genes successfully boosted the transformation efficiency of WWl50 **(Fig. 2b)**. Among the vectors carrying DR genes, pW502, which overexpresses *AtGRF5*, resulted in the highest transformation efficiency (24.73%). pW503, which overexpresses the chimeric *TaGRF4-OsGIF1* gene, and pW505, which overexpresses *ZmWUS* and *ZmBBM*, also dramatically enhanced transformation with efficiencies of 13.78% and 22.10%, respectively. The *ZmWUS* gene alone (pW504) or *ZmWUS* plus *ipt* (pW506) resulted in very weak or no increases in transformation efficiency **(Fig. 2b)**. We observed numerous growth abnormalities in plants overexpressing *ZmWUS* plus *ZmBBM*, including abnormal leaf development, expansion of the stem, and lack of an obvious stem apical meristem (pW505); however, we observed no obvious growth defects when overexpressing *AtGRF5* (pW502) or *TaGRF4-OsGIF1* (pW503) **(Fig. 2c)**. Moreover, the *AtGRF5* (pW502) overexpression transgenic watermelon plants did not exhibit any growth abnormalities after being transferred to soil **(Fig. 2d)**. Thus, we concluded that *AtGRF5* is a suitable gene to use to increase the transformation efficiency in watermelon.

### Effect of *Agrobacterium* strain and cultivar genotype on *AtGRF5*-mediated watermelon transformation

*Agrobacterium* strain dependence is a key factor affecting transformation efficiency. To identify the most suitable *Agrobacterium* strain, we introduced the pW502 vector into three *Agrobacterium* strains, GV3101, EHA105, and LBA4404. Compared with the transformation efficiency obtained using GV3101, which was used in the experiments described above (∼25%), that obtained using strain EHA105 was much lower (6.90%). Moreover, we found that LBA4404 failed to generate any positive transformation events **(Table 1)**. These results indicate that *Agrobacterium* strain selection is an important consideration for watermelon transformation, and that among the three strains we tested, GV3101 is the most suitable. This observation, together with the above results, suggests that pW502 (*AtGRF5*) delivered by strain GV3101 is a preferred strategy for watermelon transformation.

**Table 1.** The transformation efficiencies of watermelon cultivar WW150 obtained using different *Agrobacterium* strains carrying the pW502 plasmid.

| *Agrobacterium* strain | Number of explants | Positive transformants | Efficiency (%) |
| --- | --- | --- | --- |
| GV3101 | 126 | 36 | 28.12 |
| EHA105 | 116 | 8 | 6.9 |
| LBA4404 | 76 | 0 | 0 |

Many *Agrobacterium*-mediated plant transformation systems exhibit genotype dependence (Zhang *et al*., 2021). Thus, we evaluated the effect of the preferred design on transformation in another watermelon cultivar, 83166, which is the female parental line of the hybrid ‘Jingxin’. *Agrobacterium* strains GV3101, EHA105, and LBA4404 were separately used to deliver pW501 (control vector) and pW502 (*AtGRF5*) to cotyledon fragments. Unlike in the watermelon cultivar WWl50, without the help of a DR gene, the transformation efficiencies were 7.56% and 9.90% when pW501 was delivered by strains GV3101 and EHA105 respectively, suggesting that 83166 is not highly recalcitrant to transformation (**Table 2**). Compared with the other combinations, GV3101 carrying pW502 gave the highest transformation efficiency (20.72%). We also observed that LBA4404 failed to generate any positive transformation events in cultivar 83166 (**Table 2**). This result confirmed our conclusion that the *AtGRF5* gene is a robust tool that facilitates watermelon transformation, and that GV3101 is the preferred *Agrobacterium* strain to deliver these vectors.

**Table 2.** *AtGRF5*-mediated transformation of a second watermelon cultivar 83166.

| *Agrobacterium* strain | pW501 (Control) |  |  | pW502 (*AtGRF5*) |  |  |
| --- | --- | --- | --- | --- | --- | --- |
|  | Number of explants | Positive transformants | Efficiency (%) | Number of explants | Positive transformants | Efficiency (%) |
| GV3101 | 119 | 9 | 7.56 | 111 | 23 | 20.72 |
| EHA105 | 111 | 11 | 9.9 | 111 | 14 | 12.61 |
| LBA4404 | 113 | 0 | 0 | 112 | 0 | 0 |

### Application of DRs to genome editing of *ClPDS*

Targeted mutagenesis using genome editing tools such as the CRISPR/ Cas9 system is a new breeding technology that can efficiently produce the desired mutations in the target gene, and it has been applied in multiple crop species, including watermelon (Tian *et al*., 2017, Wang *et al*., 2021a, Wang *et al*., 2021b). Therefore, we tested the compatibility of AtGRF5-mediated transformation with genome editing tools in watermelon. The existence of two *Bsa*I restriction enzyme recognition sites in the original *AtGRF5* gene hampered its application in CRISPR/Cas9 mediated genome editing, as many CRISPR/Cas9 genome editing vectors are constructed using Golden Gate Assembly, which requires the use of the type IIS *Bsa*I enzyme. Thus we added the codon-optimized version of *AtGRF5*, which lacks the *Bsa*I recognition sites (**Supplemental sequences**), to the CRISPR/Cas9 genome editing plasmid pHSE401 (Xing *et al*., 2014), generating the pZHW512 vector (**Fig. 3a**). Two previously designed spacers (T1 and T2) targeting the watermelon phytoene desaturase gene *ClPDS* (Cla97C07G142100) were chosen (Tian *et al*., 2017). The resulting genome editing plasmid was transformed into WW150 cotyledon fragments. All 11 positive transgenic plants recovered were verified by Sanger sequencing. Among the 11 plants, 3 plants had mutations (#1, #5, and #9), giving a mutation efficiency of 27.30% (**Fig. 3b**). Only one mutation type, a 1-bp deletion (D1), was detected at the T1 site in #9. While at the T2 site, mutations were detected in three plants (#1, #5, and #9), with one heterozygous mutation and two biallelic mutations (**Fig. 3c-d**). As expected, the biallelic mutant exhibited evident pure albino phenotype (**Fig. 3e**). These results suggested that the DR gene *AtGRF5* could facilitate the generation of desired mutations via the CRISPR/Cas9 system.

**Fig. 3.**
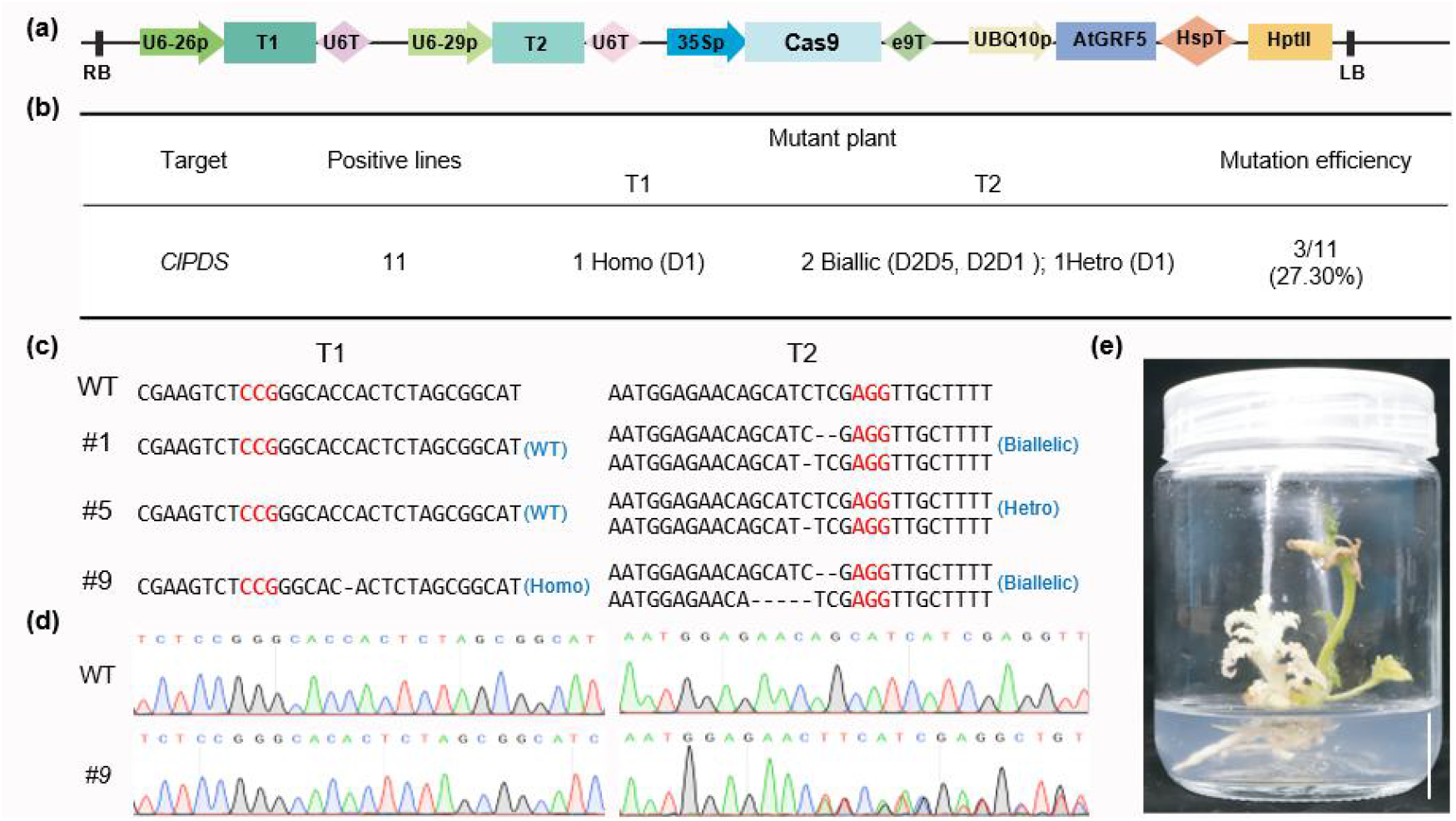
*AtGRF5* facilitates the CRISPR/Cas9-mediated genome editing of *ClPDS*. **a**. Schematic diagrams of the genome editing vector pZHW512. **b**. Summary of the *CIPDS* genome editing results. D indicates deletion. **c**. Summary of the mutation types at two target sites in the *ClPDS* gene. The PAM sequences for the two target sites are highlighted in red. **d**. Results from Sanger sequencing of the two target sites in wild type (WT) and editing line #9. **e**. Phenotype of the genome-edited *clpds* mutant plant. Scale bar = 2 cm.

## Discussion

Genetic transformation has been widely used to advance plant genome research. Although successful and highly efficient transformation systems have been established in some species, transformation remains the bottleneck in the production of transgenic and genome-edited plants for watermelon as well as many other horticultural species. Transient expression methods, such as virus-induced gene silencing using virus vectors and protoplast-based transient expression assays, have recently been established in watermelon (Igarashi *et al*., 2009, Zhao *et al*., 2016, Tian *et al*., 2017, Ren *et al*., 2018, Liu *et al*., 2020, Wang *et al*., 2021a), and these methods can be applied to study gene functions. However, stable transformation protocols are still the foundation for the genetic engineering of crop traits. Here, we have established a highly efficient transformation method for watermelon.

Fluorescent proteins and visible markers have been widely used to screen for positive transgenic events and transgene-free plants and to identify the genotype of plants (Dong *et al*., 2018, Aliaga-Franco *et al*., 2019, Yu and Zhao, 2019, He *et al*., 2020, Kong *et al*., 2021). The RUBY reporter system, which catalyzes the biosynthesis of the vividly red pigment betalain, has been established to noninvasively monitor gene expression and to screen for transformation events in both rice and *Arabidopsis* (He *et al*., 2020). Here, the fluorescent protein DsRed, which can be easily detected by a hand-held fluorescent light source, was utilized to screen for positive transformation lines, and the positive lines and negative lines could be easily distinguished.

It has been observed that overexpression of DRs improves the regeneration efficiency of recalcitrant plant species, such as wheat, maize, sorghum, and hemp (Lowe *et al*., 2016, Debernardi *et al*., 2020, Kong *et al*., 2020, Zhang *et al*., 2021). To increase the regeneration efficiency of watermelon callus, we cloned the *AtGRF5, TaGRF4-OsGIF1, ZmWUS, ZmBBM* and *ipt* genes, which were previously shown to be effective in several monocot and dicot plant species (Lowe *et al*., 2016, Debernardi *et al*., 2020, Kong *et al*., 2020), and delivered them into watermelon cotyledon fragments. We obtained the highest transformation efficiency (28.12%) when overexpressing *AtGRF5*, which 40.75-fold higher than that obtained for the empty vector control. The transformation efficiency was also dramatically increased when overexpressing *ZmWUS* plus *ZmBBM*. However, several growth defects were observed during tissue culture, similar to previous reports (Lowe *et al*., 2016, Horstman *et al*., 2017). Thus, the *ZmWUS* and *ZmBBM* genes need to be excised during tissue culture; this is often accomplished by utilizing a drought-inducible promoter to drive the expression of the recombinase Cre, which excises the DNA fragment between two LoxP sites (Lowe *et al*., 2016, Zhang *et al*., 2019). In contrast, no significant side effects were observed when overexpressing AtGRF5, similar to previous reports (Horiguchi *et al*., 2005, Kong *et al*., 2020), so no additional operation is required, which simplifies transformation and saves labor.

The CRISPR/Cas9 genome editing system has been adopted in watermelon to generate new stocks with altered sugar content, seed size, and herbicide resistance (Tian *et al*., 2017, Tian *et al*., 2018, Wang *et al*., 2021a, Wang *et al*., 2021b). However, several newly developed precise genome editing technologies, such as the base editing systems, the Prime editing system, novel CRISPR/Cas systems, and Cas9 variants recognizing a wider range of PAM sequences, that have been applied in model crops (Chen *et al*., 2019, Hua *et al*., 2019, Hua *et al*., 2020, Lu *et al*., 2021, Molla *et al*., 2021), have not been applied in watermelon. The main barrier is still the low transformation efficiency and the limited number of transformable watermelon cultivars. With the assistance of *AtGRF5*, we obtained 3 *clpds* mutant plants out of 11 transgenic plants. Co-expressing *AtGRF5* did not seem to have obvious negative effects on the genome editing efficiency. The integrated genome editing cassettes often need to be segregated out in the progeny to obtain Cas9-free plants, and this could also eliminate possible side effects when co-expressing DRs in the same genome editing vector. Moreover, overexpression of *AtGRF5* also increases the transformable cultivar range, as we obtained >20% transgenic efficiency for two watermelon cultivars, one recalcitrant and one amenable to transformation. In recent years, the addition of developmental regulators to genome editing vectors has been used more and more widely (Zhang *et al*., 2019, Debernardi *et al*., 2020, Qiu *et al*., 2021, Zhang *et al*., 2021). This strategy not only facilitates traditional genetic transformation, but also promotes the application of other precise genome modification technologies in watermelon.

In summary, we have conclusively shown that the expression of DR genes, particularly *AtGRF5*, significantly improves the transformation efficiency of watermelon without obvious negative effects on growth. Among the three *Agrobacterium* strains we tested, GV3101 was found to be the most suitable for transformation of watermelon. Using GV3101 to deliver the pW502 (*AtGRF5*) vector, we successfully achieved high transformation efficiencies of >20% for two watermelon cultivars. We believe that this strategy will facilitate the delivery of CRISPR/Cas9-based genome editing tools in watermelon, advancing gene functional studies and molecular breeding of watermelon. We think that similar strategies using DRs can also be employed to overcome the transformation barriers in many other Cucurbitaceae species.

### Experimental procedures

#### Vector construction

For the construction of pW501, the pKSE401 vector (Xing *et al*., 2014) was chosen as the backbone. The single guide RNA was removed by *Hind*III digestion and re-ligation. Then the *Cas9* gene was removed by *Xba*I and *Sac*I digestion, and replaced with DsRed by Gibson Assembly using the primer pair DsRed-F-XbaI/ DsRed-R-SacI.

For the construction of vectors with DRs, the *UBQ10* promoter and *Hsp* terminator were amplified with UBQ10p-F/UBQ10p-R and HspT-F/HspT-R, respectively, from the *Arabidopsis thaliana* Col-0 genome DNA. These fragments were introduced into the pW501 vector digested with *Hind*III by Gibson Assembly. For the expression of *WUS*, a DNA fragment containing the *Nos* promoter, *WUS* coding sequence and *CaMV* terminator was synthesized (**Supplemental sequences**). Then the *WUS* expressing cassette was amplified with the primer pair WUS-F/WUS-R and inserted into pW501 by Gibson Assembly. Other DRs were amplified with the corresponding primers and inserted into the *Kpn*I site between the *UBQ10* promoter and *Hsp* terminator.

For the coding optimization of *AtGRF5*, the codons rarely used in *Arabidopsis*, were replaced with synonymous codons with high usage frequencies. Two *Bsa*I sites in the *AtGRF5* sequence were also removed. The final sequence fragment (listed in **Supplemental sequences**) was synthesized at Sangon Biotech.

For the genome editing vector, the GRF5 expression cassette was amplified with primers ZHW512-F/ZHW512-R and inserted into the *EcoR*I site of the CRISPR/Cas9 genome editing vector pHSE401, generating the pZHW512 vector. Then two spacers targeting *ClPDS* were added by PCR with primers ClaPDS-F/ClaPDS-R using the pCBC-DT1T2 plamid(Xing *et al*., 2014) as the template, and the PCR product was inserted into the pZHW512 vector digested with *Bsa*I by Gibson Assembly.

All of the primers are listed in **Supplemental Table 1**.

#### Watermelon transformation

The genetic transformation was conducted as previously described with slight modifications (Tian *et al*., 2017). In brief, surface-sterilized watermelon seeds were sown on Murashige and Skoog (MS) solid medium for 3 days in the dark. Then the middle parts of the cotyledons without embryo were cut into 1.5 × 1.5 mm pieces as explants. *A. tumefaciens* strains harboring the indicated binary vectors were co-cultivated with the cotyledon fragments in the dark for 3 days on MS solid medium containing 1.5 mg/L 6-BA. Then the cotyledon fragments were transferred onto selective induction medium containing 2 mg/L 6-BA, 0.2 mg/L IAA, 50 mg/L Kan, and 100 mg/L Timentin. The regenerated adventitious buds were excised and transferred onto elongation medium containing 0.1 mg/L 6-BA, 0.01 mg/L NAA, 100 mg/L Timentin, and 50 mg/L Kan. Plants with full leaves and stems were transferred to rooting medium containing 1mg/L IBA and 100 mg/L Timentin. Positive transgenic events were detected using the hand-held dual-wavelength fluorescent protein excitation light source LUYOR-3415RG (Luyor Corporation, Shanghai, China) or Zeiss SteREO Discovery.V20.

#### Detection of mutations

Genomic DNA was extracted from the transgenic watermelon plants using the CTAB method. Detection of the transgene was performed with the primers Cas9-F2/Cas9-R2 and DsRed-F3/ DsRed-R3. The PCR products with the primer pair PDS-T2-F/PDS-T2-R are used as the internal control. The target regions were amplified with two pairs of primers PDS-T1-F/PDS-T1-R and PDS-T2-F/PDS-T2-R. PCR products were sent for Sanger sequencing at Sangon Biotech to determine the types of mutation. The mutation types are decoded by the DSDecodeM program (http://skl.scau.edu.cn/dsdecode/). The primers used are listed in Supplementary Table 1.

## Accession numbers

ClPDS, Cla97C07G142100.

## Acknowledgements

This work was supported by grants from Taishan Scholar Foundation of Shandong Province (tsqn202103160) and Shandong Science and Technology Innovation Funds for Huawei Zhang and Wanggen Zhang. We thank Dr. Xinping Zhang for providing watermelon seeds, and to Dr. Yun Deng for the help of transgenic plant culture in soil.

## Author contributions

H. Z. conceived this study. H.Z. and W.Z. supervised the research. W.P., Z.C., Z.H. and H.Y. performed all experiments and analyzed the data. H.Z. and W.P. wrote the manuscript with input from all authors. H.Z. and W.Z. agree to serve as the author responsible for contact and ensure communication.

## Supplementary information

**Supplemental Table 1. The list of primers used in this study**.

**Supplementary Sequences**

The complete coding sequence for different versions of *GRF5*, and the synthetic *WUS* expressing cassette.

